# Fully Bayesian longitudinal unsupervised learning for the assessment and visualization of AD heterogeneity and progression

**DOI:** 10.1101/854356

**Authors:** Konstantinos Poulakis, Daniel Ferreira, Joana B. Pereira, Örjan Smedby, Prashanthi Vemuri, Eric Westman, for the Alzheimer’s Disease Neuroimaging Initiative

**Affiliations:** Division of Clinical Geriatrics, Department of Neurobiology, Care Sciences and Society, Karolinska Institutet, Stockholm, Sweden; Department of Biomedical Engineering and Health Systems (MTH), KTH Royal Institute of Technology, Stockholm, Sweden; Department of Radiology, Mayo Clinic, Rochester, MN, USA; Department of Neuroimaging, Centre for Neuroimaging Sciences, Institute of Psychiatry, Psychology and Neuroscience, King’s College London, London, UK

**Keywords:** Alzheimer’s disease, brain atrophy, neuroimaging, atrophy progression, longitudinal cluster analysis

## Abstract

Tau pathology and regional brain atrophy are the closest correlate of cognitive decline in Alzheimer’s disease (AD). Understanding heterogeneity and longitudinal progression of brain atrophy during the disease course will play a key role in understanding AD pathogenesis. We propose a framework for longitudinal clustering that: 1) incorporates whole brain data, 2) leverages unequal visits per individual, 3) compares clusters with a control group, 4) allows to study confounding effects, 5) provides clusters visualization, 6) measures clustering uncertainty, all these simultaneously. We used amyloid-β positive AD and negative healthy subjects, three longitudinal sMRI scans (cortical thickness and subcortical volume) over two years. We found 3 distinct longitudinal AD brain atrophy patterns: a typical diffuse pattern (n=34, 47.2%), and 2 atypical patterns: Minimal atrophy (n=23 31.9%) and Hippocampal sparing (n=9, 12.5%). We also identified outliers (n=3, 4.2%) and observations with uncertain classification (n=3, 4.2%). The clusters differed not only in regional distributions of atrophy at baseline, but also longitudinal atrophy progression, age at AD onset, and cognitive decline. A framework for the longitudinal assessment of variability in cohorts with several neuroimaging measures was successfully developed. We believe this framework may aid in disentangling distinct subtypes of AD from disease staging.

## 1. Introduction

Imaging biomarkers of brain morphology have been increasingly used in research and clinical routine during the last decades [1]. More specifically, dementia research has utilized such markers for the investigation of disease-related patterns from populations around the world and many cohorts with complete neuroimaging data are now available to the research community [2,3]. Structural neuroimaging markers are also used for selection of participants for clinical trials in Alzheimer’s disease (AD) [4]. The availability of longitudinal data provides us with the opportunity to assess changes over time in healthy and pathological individuals. A new challenge for the imaging research community is the incorporation of longitudinal information in their study designs [5]. Some attempts to utilize those data in order to understand disease progression include the EuroPOND and the TADPOLE projects^1^. Challenges other than longitudinal information include the assessment and fixation (*ceteris paribus*) of different study effects, the meaningful visualization of group differences and finally the simultaneous optimization of all these procedures for the sake of reproducibility in the presence of pragmatic sample sizes.

Unsupervised classification (clustering) is widely applied to neuroimaging data when the aim is to unveil heterogeneous features within samples [3]. When samples include only one diagnosis, a common use of clustering methods is to investigate whether the neuroimaging measures of interest show heterogeneous patterns within that same diagnostic label. Several studies have investigated the heterogeneity in AD with the aim to define disease specific subtypes [6,7,7-13]. When samples include more than one diagnosis, the main aim of unsupervised clustering methods is to investigate whether neuroimaging markers can be used to distinguish between the diagnostic classes without specifying them with a label. The clustering methods that are used today are mostly cross-sectional, in the sense that they utilize baseline observations for a set of individuals. In the AD research field, many studies have focused on the unbiased identification of cortical and subcortical patterns of atrophy with structural MRI (sMRI). One recent study utilizes longitudinal atrophy markers to find sets of brain regions with common progression patterns [14], However, to date no cluster-based study has included longitudinal atrophy data in their method scheme, in order to identify groups of individuals with similar atrophy trajectories and our current study intends to meet this necessity.

In studies where the aim is to investigate neuroimaging measures in association to some clinical outcome, we often wish to account for or exclude the effect of confounders that can potentially introduce bias and may drive the results of our analysis. More specifically, in connection with cluster analysis, two approaches are widely used in the literature. The first approach, is called the residual (de-trending) method [15,16]. In this approach a “correction” is applied to a neuroimaging measurement with respect to a confounder that should not affect the results of the main analysis. The adjusted measurement will not be correlated with the confounder anymore. After that, one applies the clustering algorithm on the de-trended data [10,11,17-19]. When using the de-trending approach, the statistical tests that we need increase dramatically in numbers (one correction for each vertex/voxel/region of interest). Moreover, the cluster parameters are not optimized conditional to the original data but given the artificial data (de-trended data). All these features can make the interpretation of results more difficult and introduce errors in reproducibility, since the results are based on a chain of statistical procedures that are not connected in statistical terms. According to the second approach (for confounders in a clustering study), it is suggested to incorporate the effect that we want to account for in the analysis [9,20]. This can be achieved with the addition of one fixed effect in the case of a statistical clustering model.

Another important feature of a neuroimaging clustering study is the comparison of differences in brain morphology between clusters of individuals. This comparison commonly involves, i) groups of the same pathology with different atrophy patterns or ii) a pathological group and a cognitively unimpaired group with similar demographical characteristics, or other combinations of comparisons between groups. This step is either incorporated in the clustering procedure, or it is performed as an independent post-clustering step. When this step is not included in the clustering procedure, but added as a separate step, we need to correct the resulting images for multiple statistical comparisons since multiple models are implemented for that purpose. This issue can be avoided in the case of a simultaneous clustering and visualization.

Previous clustering studies grouped AD patients based on sMRI features from a single time-point [8,10,13,17,20-22]. Their conclusions were based on a single observation in time and the chance that those clusters reflect different stages of the disease and not particular patterns of atrophy (distinct AD subtypes) cannot be excluded. A longitudinal clustering design can reduce the risk that the results will reflect different disease stages. Even if the clusters reflect different disease stages, we can infer them with higher certainty than in a cross-sectional study. Moreover, the follow up MR acquisitions can be irregularly distributed between subjects, may drop out or miss certain visits. We model this feature in order to obtain accurate estimates of atrophy progression.

In this study, we aimed to design and assess a framework for longitudinal clustering that incorporates: 1) simultaneous clustering of several longitudinal neuroimaging measures (multivariate data over time), 2) information for individuals with irregularly sampled observations, 3) comparison of the clusters with a control group, 4) the study and fixation (optional) of effects that should not drive the resulting clusters, 5) visualization of the resulting clusters for interpretation, 6) measures of uncertainty in the clustering. Our overall goal is to perform all the aforementioned methodological steps in one statistical model in order to avoid the statistical pitfalls of a “pipeline” study that limits the ability to correctly identify disease mechanisms because of weak statistical inferences. The designed framework is applied to longitudinal sMRI data of mainly amyloid-β (Aβ) positive AD patients and Aβ negative cognitively unimpaired (CU) subjects over a period of two years (three sMRI time points). To assess the results from our new longitudinal clustering framework, we included all data with longitudinal information from our previous cross-sectional clustering study [13]. This allows us to compare the results from cross-sectional and longitudinal clustering in the same dataset. To be able to estimate cluster-specific atrophy trajectories in time is a very important aspect that has been overlooked by previous cross-sectional AD subtypes studies [23]. This approach will provide relevant data in order to answer an unresolved question in the field, i.e., whether “AD subtypes” are truly distinct subtypes or they are rather stages of the AD pathophysiological process reflected in crosssectional analyses.

## 2. Results

### 2.1 Clustering evaluation

The reported results are based on 750000 iterations with 500 iterations thinning where 250000 iterations were burn-in period, which therefore saved 1000 MCMC samples. The distributions of the estimated parameters started converging after the burn-in samples and it remained stable afterwards for the rest of the simulations. As expected, the general tendency of the deviance for the different models decreased with increasing amount of clusters (Supplementary table 1). The different initializations brought various outputs from which the one with the packages’ default settings was the worst in terms of deviance. The model with initialization in the means of the clusters from our previous study [13] and the addition of uniform noise for 8 clusters was optimal in terms of quality.

Figure 2 shows the multidimensional scaling coordinates of the component-subject probability matrix. Subjects are coloured dependent on the cluster that they belong to. Clusters 7 and 8 comprised 2 subjects each (Figure 2, A) and were thus considered outlier clusters under the maximum probability rule. Moreover, the classification of subjects into clusters with high posterior density (HPD) intervals showed that 3 out of 72 subjects (1 subject from cluster 7 and 2 subjects from cluster 2) had uncertain classification (Figure 2, B). These subjects were excluded from the post hoc analysis and interpretation. The data of the 6 subjects (outlier clusters 7 and 8, and HPD interval uncertain classified subjects; one of the subjects belonged in outlier cluster 7 and had uncertain classification under the HPD intervals method too) are presented in Supplementary figure and table 3).

**Figure 1.**
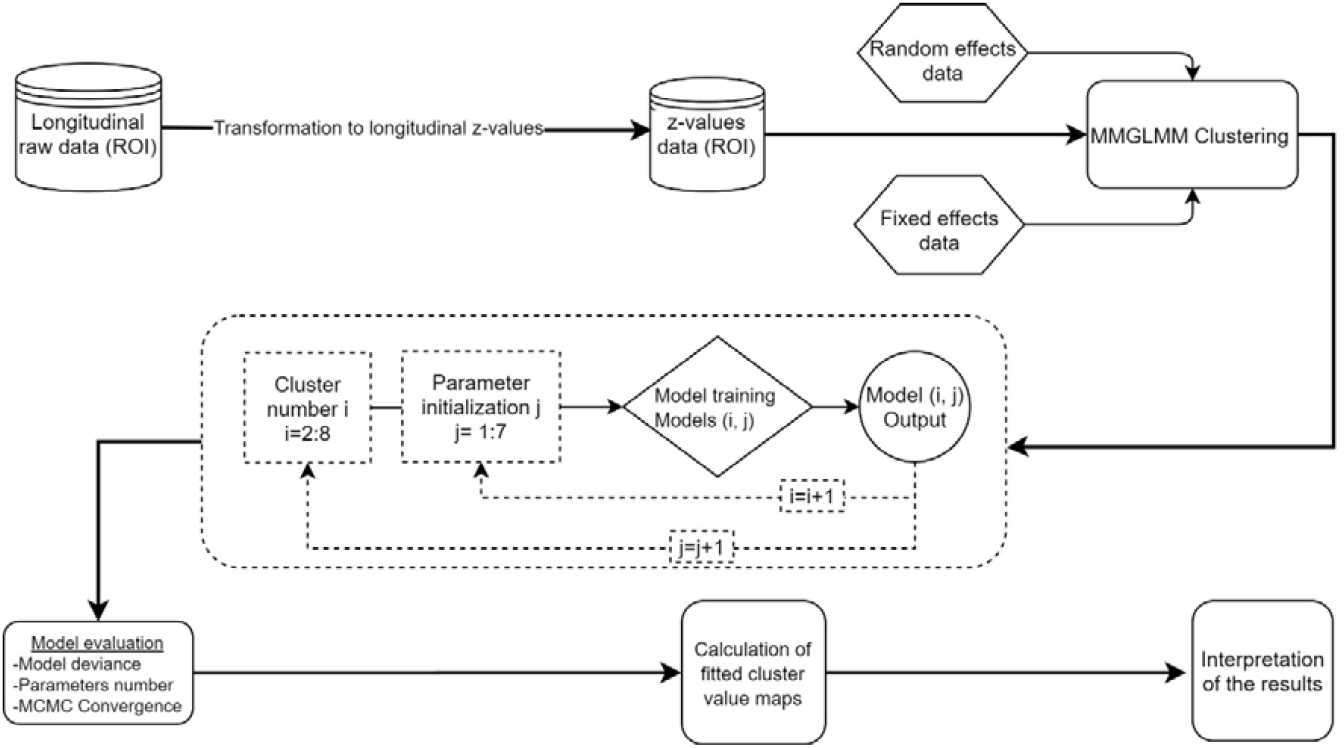
Flowchart of the analysis. The schematic representation of the analysis shows that all the steps after the data standardization are accomplished within the clustering and not in separate pipeline fashion like steps. ROI: region of interest, MMGLMM: Multivariate Mixture of Generalized Mixed effect Models.

**Figure 2.**
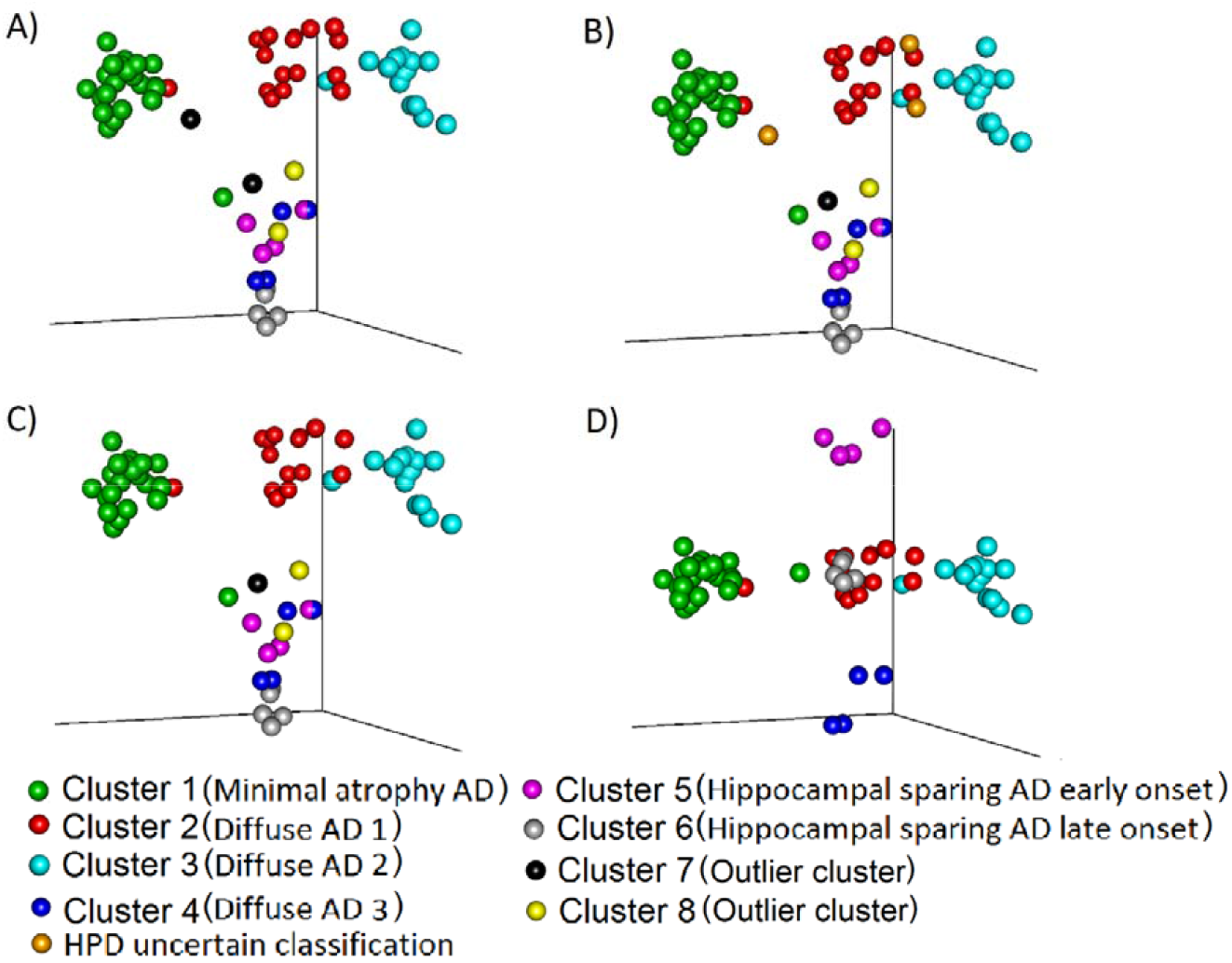
Comparison of maximum probability and HPD interval classifications. Three-dimensional representation of (Multidimensional scaled (MDS)) component-individual probabilities matrix (This matrix includes the probability of each subject to be in any of the clusters). The scatter plots represent subjects and are coloured according to the clustering based on two approaches, maximum probability and highest posterior density intervals (HPD). A) Subjects are coloured based on maximum probability classification (MDS components 1, 2 and 3). B) Subjects are coloured based on HPD intervals classification. In comparison to A, in B we added the uncertain classification with orange colour (Two subject from cluster 2 and 1 subject from cluster 7 cannot be classified to any cluster with high certainty). C) Colours are the same as in B, but we excluded from the plot the HPD uncertain classification subjects: orange and the outlier clusters 7: black and 8: yellow. D) The subjects are coloured exactly as in C but the MDS components 1, 2 and 5 are plotted, to showcase the separation between Cluster 4, 5 6. The names in the parenthesis after the cluster numbers refer to the figure 3 and table 2.

The remaining 66 subjects were used for further analysis. The separation between the 6 clusters in terms of how probable it is for their subjects to belong to the same cluster is seen in figure 2 C where the clusters 1, 2 and 3 are clearly separated from each other. An additional visualization of the 1^st^, 2^nd^ and 5^th^ multidimensional scaled (MDS) components shows the separation between Cluster 4, 5 and 6 (Figure 2, D).

### 2.2 Cluster characterization

Three main patterns of atrophy were found in the dataset: i) typical AD pattern (clusters diffuse 1, 2 and 3) (Figure 3 B), ii) a minimal atrophy pattern (Figure 3A) and iii) a hippocampal sparing pattern (hippocampal sparing early and late onset) (Figure 3C).

**Figure 3.**
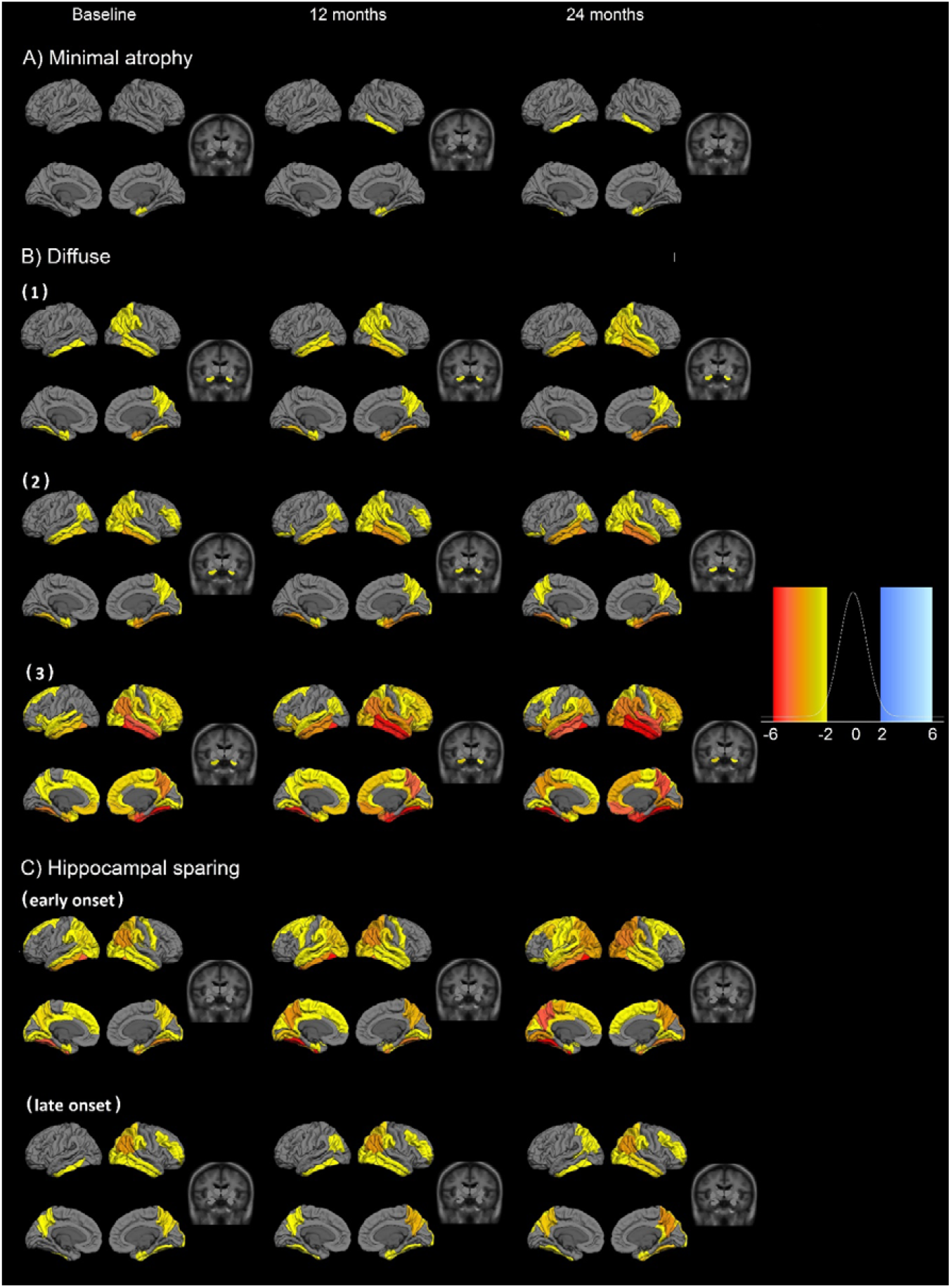
Fitted values for cortical thickness and subcortical volumes for the different patterns of atrophy. Atrophy fitted values of the 6 longitudinal atrophy patterns for the AD sample. Each row presents the median fitted values of the cortical and subcortical atrophy of the 6 components for three time points (baseline, 12 and 24 months from the first measurement). The data are presented as cognitively unimpaired group z-scores. A: Minimal atrophy pattern, B: Diffuse AD atrophy pattern, C: hippocampal sparing AD atrophy pattern. Fixed effects: Intracranial volume = average Intracranial volume, Sex= female, Age = 75 years, Time from onset of dementia = 5 years, Education = 16 years, CSF Aβ1-42 = 100 pg/ml, CSF Ptau181P = 50 pg/ml. Data are presented as standard deviations below the estimated mean of the healthy cognitively unimpaired population.

The Minimal atrophy cluster is characterized by initial atrophy in the entorhinal cortex (right) and longitudinal decrease in thickness first in the right and then in the left inferior temporal gyrus during the 24 months of follow up (Figure 3A). The atrophy patterns in the three Diffuse clusters (reported as typical AD), more closely follow the NFTs pattern suggested by [24], However, differences do exist and may be attributed to age (even after correcting for this effect). Further, the atrophy in the Diffuse 3 cluster is more advanced (Figure 3B) and these subjects have lower cognitive performance (Table 2). This may be the reason for why the Diffuse 3 cluster only had MRI for baseline and 12 months follow-up. Within the hippocampal sparing AD subtype two clusters are observed. The degree of atrophy as well as the age at onset of dementia differentiate these two clusters (Figure 3C, Table 2). The early onset hippocampal sparing cluster has greater level of atrophy at baseline and accumulates atrophy faster over time, in contrast to the late onset hippocampal sparing cluster. In both clusters the precuneus and the inferior parietal gyri (Figure 3C) are atrophied. For a more comprehensive understanding of the atrophy distributions in the cortex of the different clusters, we can also utilize the 1^st^ and 3^rd^ quartile images that present the dispersion around the mean cortical atrophy of each cluster (Supplementary Figure 1).

**Table 1.** Sample demographics.

|  |  |  |
| --- | --- | --- |
| **N** | 72 | 31 |
| **Females N (%)** | 34 (47.2%) | 15 (48.4%) |
| **Age mean (sd)** | 76 (7.4) | 74 (4.4) |
| **Age at disease onset median(mad)** | 71 (8.9) | - |
| **Years of education median(mad)** | 16 (3) | 16 (3) |
| **MMSE median(mad)** | 24 (1.5) | 29 (0) |
| **CDR global score median(mad)** | 0.72 (0.25) | 0 (0) |
| **ApoE e4 allele carrier N (%)** | 50 (69.4%) | 3 (9.7%) |
| **CSF Aβ<sub>1-42</sub>, median(mad)** | 137.38 (23.98) | 234.11 (20.88) |
| **CSF pTau<sub>181</sub>, median(mad)** | 37.5 (12.6) | 18 (4.45) |
| **ADAS word recall mean (sd)** | 6.17 (1.43) | 2.81 (0.95) |

Mad: maximum median distance, MMSE: mini mental state examination, ADAS: Alzheimer's disease assessment scale, CDR: Clinical Dementia Rating, CSF: cerebrospinal fluid, ADAS: Alzheimer's disease assessment scale. CSF values are in pg/ml. CU: cognitively unimpaired.

**Table 2.** Demographic and clinical characteristics of the clusters

|  | Minimal atrophy |  | Diffuse 1 (Typical AD) |  | Diffuse 2 (Typical AD) |  | Diffuse 3 (Typical AD) |  | Hippocampal sparing early onset |  | Hippocampal sparing late onset |  |
| --- | --- | --- | --- | --- | --- | --- | --- | --- | --- | --- | --- | --- |
|  | m0 | m24 | m0 | m24 | m0 | m24 | m0 | m12 | m0 | m24 | m0 | m24 |
| Demographics |  |  |  |  |  |  |  |  |  |  |  |  |
| N (%) | 23 (35%) | 20 | 15 (23%) | 13 | 15 (23%) | 12 | 4 (6%) | 4 (6%) | 4 (6%) | 4 (6%) | 5 (7%) | 2 |
| Females N(%) | 9 (39.1%) | 9 (45%) | 7 (46.7%) | 6 (46.2%) | 8 (53.3%) | 6 (50%) | 2 (50%) | 2 (50%) | 2 (50%) | 2 (50%) | 2 (40%) | 1 (50%) |
| Age | 76 (10.4) | 78 (12.6) | 74 (5.9) | 76 (5.9) | 79 (3) | 81 (3.7) | 73.5 (3.7) | 73.5 (3.7) | 66.5 (6.7) | 66.5 (6.7) | 70 (14.8) | 74.5 (23) |
| Age disease onset | 70 (10.4) | 71.5 (11.9) | 68 (7.4) | 72 (7.4) | 75 (4.4) | 73.5 (5.2) | 68.5 (5.2) | 68.5 (5.2) | 59.5 (3.7) | 59.5 (3.7) | 67 (11.9) | 70 (23.7) |
| Years of education | 16 (4.4) | 15 (3.7) | 14 (3) | 14 (3) | 12 (5.9) | 13.5 (3.7) | 16 (3) | 16 (3) | 18 (0) | 18 (0) | 16 (3) | 15 (1.5) |
| Apoe e4 allele carrier N(%) | 18 (78.3%) | 15 (75%) | 12 (80%) | 10 (76.9%) | 9 (60%) | 8 (66.7%) | 3 (75%) | 3 (75%) | 2 (50%) | 2 (50%) | 2 (40%) | 1 (50%) |
| CSF biomarkers |  |  |  |  |  |  |  |  |  |  |  |  |
| Aβ1-42 | 129.81 (17.57) | 129.38 (16.54) | 136.83 (41.35) | 140.09 (27.1) | 140.37 (25.43) | 153.81 (11.4) | 153.81 (11.4) | 153.81 (11.4) | 128.8 (17.09) | 128.8 (17.09) | 143.32 (5.69) | 145.71 (9.24) |
| PTAU181P | 38 (13.34) | 37.5 (14.83) | 44 (20.76) | 44 (20.76) | 35 (5.93) | 35.5 (5.93) | 35.5 (5.93) | 35.5 (5.93) | 36.5 (14.08) | 36.5 (14.08) | 50 (35.58) | 36 (20.76) |
| Cognitive measures |  |  |  |  |  |  |  |  |  |  |  |  |
|  | Median (mad) | Annual change (se) | Median (mad) | Annual change (se) | Median (mad) | Annual change (se) | Median (mad) | Annual change (se) | Median (mad) | Annual change (se) | Median (mad) | Annual change (se) |
| MMSE | 25 (1.5) | -0.8(0) | 22 (1.5) | -2.2(0.1) | 24 (1.5) | -1.9(0) | 21.5 (0.7) | -3.3(0.1) | 25.5 (0.7) | -4.1(0.1) | 23 (1.5) | -2.6(0.1) |
| CDR global | 0.65 (0.24) | 0.15(0) | 0.77 (0.26) | 0.27(0.01) | 0.7 (0.25) | 0.23(0.01) | 0.88 (0.25) | 0.25(0.02) | 0.75 (0.29) | 0.37(0.01) | 0.6 (0.22) | 0.22(0.01) |
| ADAS 11 |  |  |  |  |  |  |  |  |  |  |  |  |
| ADAS Q1 Word recall | 5.3 (1) | 0.5(0) | 7 (1.5) | 0.2(0) | 6.3 (1) | 0.4(0) | 8.3 (1) | 0.7(0) | 6.7 (2.7) | 0.7(0.1) | 6.3 (1) | 0.5(0) |
| ADAS Q2 Commands | 0 (0) | -0.1(0) | 0 (0) | 0.2(0) | 0 (0) | 0.1(0) | 1 (0) | 0.3(0) | 0 (0) | 0.6(0) | 0 (0) | 0.2(0) |
| ADAS Q3 Constructional praxis | 1 (0) | 0.1(0) | 1 (0) | 0.1(0) | 1 (0) | -0.1(0) | 1.5 (0.7) | 0.3(0) | 1 (0) | 0.6(0) | 1 (0) | 0.2(0) |
| ADAS Q4 Delayed word recall | 8 (1.5) | 0.5(0) | 10 (0) | 0(0) | 9 (1.5) | 0.4(0) | 10 (0) | 0(0) | 9 (1.5) | 0.3(0) | 8 (0) | 1.2(0) |
| ADAS Q5 Naming objects and fingers | 0 (0) | 0.1(0) | 1 (0) | 0.4(0) | 0 (0) | 0.3(0) | 0.5 (0.7) | 1.2(0) | 0.5 (0.7) | 0.4(0) | 0 (0) | 0(0) |
| ADAS Q6 Ideational praxis | 0 (0) | 0.2(0) | 0 (0) | 0.6(0) | 0 (0) | 0.2(0) | 0 (0) | 0.7(0) | 0.5 (0.7) | 0.7(0) | 0 (0) | 0.4(0) |
| ADAS Q7 Orientation | 1 (1.5) | 0.4(0) | 2 (1.5) | 1.1(0) | 2 (1.5) | 1.2(0) | 3.5 (1.5) | 0.8(0.1) | 1 (0) | 2(0) | 2 (1.5) | 1.1(0) |
| ADAS Q8 Word recognition | 6 (3) | 0.6(0) | 7 (3) | 1.2(0) | 8 (3) | 0.4(0) | 11 (0.7) | -0.4(0.1) | 7.5 (4.4) | 0.9(0.1) | 4 (1.5) | 2.7(0) |
| ADAS Q9 Remembering test instructions | 0 (0) | 0.1(0) | 0 (0) | 0.3(0) | 0 (0) | 0.1(0) | 0 (0) | 0.7(0.1) | 0 (0) | 0.9(0) | 0 (0) | -0.2(0) |
| ADAS Q10 Language | 0 (0) | 0.1(0) | 0 (0) | 0.2(0) | 0 (0) | 0.2(0) | 0 (0) | 0.2(0) | 0 (0) | -0.1(0) | 0 (0) | 0.3(0) |
| ADAS Q11 Word finding difficulty | 0 (0) | 0.4(0) | 0 (0) | 0.5(0) | 0 (0) | 0.4(0) | 1.5 (0.7) | 0.8(0.1) | 1.5 (1.5) | 0.1(0.1) | 0 (0) | 0.5(0) |
| ADAS Q12 Comprehension of spoken language | 0 (0) 0.1(0) | 0.1(0) | 0 (0) | 0.1(0) | 0 (0) | 0.1(0) | 0 (0) | 1.3(0.1) | 0 (0) | -0.4(0) | 0 (0) | 0.1(0) |

The data are presented as median (median absolute distance) unless otherwise stated. CSF: cerebrospinal fluid, MMSE: mini mental state examination, CDR: Clinical Dementia Rating, ADAS: Alzheimer's disease assessment scale, m0 = first visit, m24 = visit after 24 months. Annual changes in the cognitive assessment scales were estimated with linear regression (follow up data as predictor, two parameters and variance estimation). Standard errors of the estimated parameters are included in brackets. No statistical tests between groups are performed due of small sample sizes in some of the groups. CSF values are in pg/ml.

The six clusters did not differ in terms of sex distribution but they differed in the years of formal education, with the average education being around 16 years (Table 2). The lowest and highest median years of education are observed in the Diffuse 2 cluster (12 years) and the Hippocampal sparing early onset (18 years). The two clusters with hippocampal sparing patterns of atrophy differ in several aspects such as the age of onset of dementia. The Minimal atrophy cluster has the slowest decline over time in MMSE and CDR, while the Hippocampal sparing early onset cluster has the steepest decline (Table 2, Supplementary figure 4). The Hippocampal sparing early onset cluster also has a steep decline in constructional praxis, and the greatest deficits in ideational praxis at baseline, but not steeper than the Diffuse 3 group. Although the Minimal atrophy cluster has the best scores in all the ADAS subscales at baseline, the Hippocampal sparing late onset group has a better score in the word recognition task at baseline but declines very fast during the next two years (2.7 points/year).

### 2.3 Comparison to previous results

The atrophy patterns of the different clusters have similarities with our previously reported cross-sectional results on AD subtypes (Poulakis et al. 2018), while the differences stem from the longitudinal information that is now added in the algorithm.

The subjects in the cross-sectional study [13] that were assigned to the Diffuse 1 subtype are now distributed in more than one cluster with the highest prevalence in the Diffuse 1 and 2 clusters (Table 3). Three subjects from the cross-sectional Diffuse 2 cluster are now in the Diffuse 3 (2 subjects) and cluster 8 (1 subject, outlier cluster). All the seven subjects from the cross-sectional hippocampal sparing subtype are still in the Hippocampal sparing clusters. Three subjects, assigned to the limbic predominant atrophy pattern in the cross-sectional study are now in cluster 7 (outlier cluster), Diffuse 1 and the HPD group. The subjects in the minimal atrophy group are still mainly in Minimal atrophy in the present study (17 subjects out of 20) while two subjects are assigned to the Diffuse 1 cluster and one subject to the Hippocampal sparing late onset cluster. Out of four CSF Aβ_1-42_ negative AD subjects that are included in the current study, one subject is assigned to the longitudinal diffuse 2 cluster (was in the cross-sectional diffuse 1 cluster), one in the longitudinal outlier cluster 7 (was in the cross-sectional limbic predominant cluster) and two are assigned to the longitudinal minimal atrophy cluster (both subjects were in the cross-sectional minimal atrophy cluster) (Table 3).

**Table 3.** Correspondence matrix

|  | Names of clusters | Longitudinal clustering results |  |  |  |  |  |  |  |  | Sum |
| --- | --- | --- | --- | --- | --- | --- | --- | --- | --- | --- | --- |
|  |  | Minimal Atrophy | Diffuse 1 (Typical AD) | Diffuse 2 (Typical AD) | Diffuse 3 (Typical AD) | Hippocampal sparing early onset | Hippocampal sparing late onset | Cluster 7 | Cluster 8 | HPD uncertain |  |
| Cross-sectional clustering results | Diffuse 1 (Typical AD) | 6 | 12 | 15 | 2 | 1 | 0 | 0 | 1 | 2 | 39 |
|  | Diffuse 2 (Typical AD) | 0 | 0 | 0 | 2 | 0 | 0 | 0 | 1 | 0 | 3 |
|  | Hippocampal sparing | 0 | 0 | 0 | 0 | 3 | 4 | 0 | 0 | 0 | 7 |
|  | Limbic predominant | 0 | 1 | 0 | 0 | 0 | 0 | 1 | 0 | 1 | 3 |
|  | Minimal atrophy | 17 | 2 | 0 | 0 | 0 | 1 | 0 | 0 | 0 | 20 |
|  | Sum | 23 | 15 | 15 | 4 | 4 | 5 | 1 | 2 | 3 | 72 |

Correspondence between the assignments of subjects in the cross sectional clustering [13] and the current longitudinal study (clustering according to the highest posterior density intervals). The cross-sectional study clusters are in the rows and the longitudinal study clusters are in columns.

## 3. Discussion

The optimization of the longitudinal clustering model provided us with interesting findings that support its future use in imaging research for studying heterogeneity in healthy and pathological ageing. Clustering with several longitudinal measures that were irregularly sampled was successfully achieved. We incorporated information from a cognitively unimpaired sample to calculate age-corrected levels of atrophy, while avoiding the need to correct for multiple comparisons. This allowed the direct visualization of atrophy trajectories. Estimated subject-component probabilities made it possible to assess whether subjects are clustered with high certainty or not. All these features provided us with useful insights that substantially helped in the interpretation of the clusters. Moreover, the assessment of some study effects within the model can potentially assist to investigate which brain regions are statistically associated with them. The framework identified and characterized three overall groups of AD subjects with distinct atrophy patterns with different trajectories over time and cognitive profiles.

### 3.1 Longitudinal clustering initialization and performance

The use of the current dataset helped the evaluation of our framework, because the patterns of atrophy at baseline are known from our previous results [13]. Thus, the longitudinal information incorporated in our framework helped us study if AD atrophy patterns at baseline change when information about the course of the disease is added. The optimization process was longer and more intensive for larger numbers of clusters, since every additional component increased the number of new parameters to be estimated. Initially, the packages’ default values for the parameters were used to see the extent of adaptation of the model to the data without any help of locally optimal solutions. The results showed that the model tends to produce 1-2 components that represent the actual dataset, while the rest of the components have non-sensible values. Moreover, the subjects were classified with high certainty in these 1-2 realistic components. This is advantageous because it means that the probabilistic clustering correctly identifies the components that represent the data in the best way. However, the rest of the components remained empty, which is a sign that the algorithm estimates components with zero presence in the dataset if it is not given some hints on where the data actually lie in the parameter space. The model with default initial values was not considered adequate to describe the dataset since too many parameters had no meaning in our application.

The decision to start the algorithm from the cross-sectional clustering results showed that when the algorithm is fed with initial information for the mixed-effect parameters, the components are more meaningful, in the sense that if not all, almost all the components have some subjects in them. However, some of the cross-sectional solutions may not be optimal since they were not specifically adjusted to the dataset. For example, when we started the cluster intercepts optimization (initial values) from a k-means cross-sectional clustering solution, the resulting model had low quality, because a more sophisticated method is needed to find suitable clusters that can describe the AD dataset. In contrast, when we started the optimization with initial values (for the cluster intercepts) from the cross-sectional AD subtypes results [13], the model received the best quality scores among the different initializations. The lack of initial values for the slopes of each cluster (we set the initial slopes to zero due to lack of longitudinal cluster information) might be the reason behind the superiority of a solution with the introduction of uniform noise. In this way, we let the algorithm search for an optimal solution that may not fit (in the parametric space) exactly to the previous study’s solution but in a parametric region close to it. Thus, we give more flexibility to the optimizer of the model to end up in the same values (as the cross-sectional study), only if these are the optimal ones. In this way, we avoid stumbling on a local optimum.

We also checked that the variance of the posterior distribution of the fixed effects was considerably smaller than the large prior value that we set it to, in accordance to the Supplementary material of the paper where the clustering methodology was presented [26]. The cluster-specific parameters (random effects) such as the mean, covariance matrix and proportion of cluster parameters were the most demanding parameters to optimize, especially in the case of 7 and 8 cluster solutions. The visual inspection of the MCMC trace plots for these parameters showed large steps at the first thousand iterations (burn in period and some iterations later) and then a stable distribution (good chain mixing) is produced.

The idea behind calculating a composite measure of model quality was inspired by the fact that all chains converged perfectly for none of the models. However, some autocorrelation was allowed to exist, which often happens in applications of Bayesian statistics (Gelman et al. 2013, chapter 11). We accepted a certain extent of autocorrelation within chains but did not accept any solution with high values (Dobson and Barnett 2018, chapter 13). The number of chains that had some autocorrelation among the random effects of the selected model was only 6% of the overall parameters, which is a reasonable amount (considering that the chains are generally mixing sufficiently well). Criteria such as Akaike’s information criterion and Bayesian information criterion provide information about deviance, parameter number and sample size, but disregard uncertainty in the model parameters (Bishop, 2007 page 33). In our approach, we used information about uncertainty and quality of Bayesian optimisation together with the deviance, to exploit the quality of the deeper features of our model structure.

This proposed clustering provides us with two additional types of information apart from the cluster assignment: 1) which subjects in a cohort are not well represented by one cluster (i.e. outliers), 2) which subjects are at risk of shifting from one cluster to another (for example from a cognitive normal cluster to a pathological cluster, i.e. HPD uncertain). In this study we also decided that two clusters of the output model should be considered as outliers. The number of observations that are needed in order to treat a cluster as an outlier is not well defined in the literature. However, we decided that 2 subjects are too few to allow an interpretation of the cluster characteristics and/or an extrapolation to the AD population. The estimated components should have a certain presence in the population in order to interpret them; otherwise the weakness of these clusters might introduce noise in the understanding of heterogeneity in the context of this application. The first outlier cluster includes one subject who is characterised by little bilateral temporal atrophy as well as subtle right hippocampal atrophy at 12 months, that cannot be captured at the 24-months observation. This may be a matter of longitudinal data preprocessing deviance in the volume estimation. The second outlier cluster has two subjects with typical AD cortical atrophy. However, one of them has no subcortical atrophy and the other has subtle left hippocampal atrophy that cannot be captured at the 12-moths assessment, together with large bilateral caudate volumes (in all timepoints) in comparison to the CU sample. For the sake of transparency, the data of the subjects that were excluded from interpretation are reported in the Supplementary Figure 3 and Supplementary Table 3. Overall, the longitudinal clustering model combined with a priori chosen initial values for the cluster specific parameters produced reasonable cluster estimates for meaningful interpretation of our longitudinal neuroimaging data.

### 3.2 AD subtypes and longitudinal clustering of brain atrophy

The results of the model support that information about atrophy trajectories has the potential to advance our current understanding about the heterogeneity within AD. Since we estimated trajectories of atrophy over time for each individual, we have more complete information to answer whether AD subtypes reported in cross-sectional studies are a result of different disease stages or they are actually distinct subtypes. We identified three main patterns of atrophy with different atrophy signature over time: i) a typical AD pattern, ii) an AD pattern where the cortex is mainly involved while the hippocampus is relatively spared and iii) a minimal atrophy pattern were subjects exhibited mild or no atrophy in cortical and subcortical regions. Within typical AD, we found three atrophy patterns. The most typical AD like atrophy pattern is observed in the Diffuse 1 cluster that has all the demographical and cognitive characteristics of AD, such as the age of AD onset (>65 years of age), MMSE (18.5±7.1) and CDR global (1.3±0.8) [6,7,22,30]. The Diffuse 2 cluster is not substantially different in median fitted values from the Diffuse 1 cluster. However, the higher age at onset (7 years older) and the percentage of females (53.5% in comparison to 46.7%) in the Diffuse 2, together with the atrophy distribution dispersion in this cluster provided by the 1^st^ and 3^rd^ quartile atrophy maps (supplementary figure 1), are somehow reminiscent of the AD subtype known as limbic predominant AD [6,7,13,17,22,31]. We speculate, given the longitudinal data and the previous cross-sectional study results [13], that the limbic predominant atrophy patterns is part of the AD disease staging rather than a distinct subtype. For some reason this cluster has later onset, but patients seem to follow the Braak staging for NFT distribution and spread, hence they will likely develop typical AD at advanced stages. Regarding the Diffuse 3 cluster, this is the most atrophied group of subjects in this dataset, its cognitive scores are very low and its frequency in the data is very small (4 subjects). Being already reported in previous results of our group [13], we can now show that this group consists of subjects with already advanced atrophy at the time of the MRI. The model estimates a random intercept for each ROI at the time of the first MRI acquisition for each subject. Therefore, the few subjects of the diffuse 3 cluster were separated from the other two diffuse atrophy clusters, since they had very low intercepts (great amount of atrophy) in the limbic areas and association cortex, as we can see in Figure 3.

The Minimal atrophy cluster, that includes subjects with minimal atrophy changes over time, is a cluster of considerable interest since the low amount of atrophy correlates well to the very slow cognitive decline in this group. The frequency of minimal atrophy in the current study is higher than in previous studies [7,13,22], most probably due to the longitudinal design that allows subjects with slow cognitive decline to be followed up for a longer period. This interpretation is supported by the finding that the Diffuse 3 cluster, the more severe group, is the only cluster that did not have 24 months follow up (early drop-out). It is proposed that tau-related pathophysiology and abnormal levels of Aβ alone without significant atrophy are enough to produce the dementia symptoms in the minimal atrophy subtype [22], perhaps through disruption of relevant brain networks in the absence of overt brain atrophy (Ferreira et al 2019), in the context of lower cognitive reserve [25,32].

The hippocampal sparing subtype with accumulation of atrophy mainly in cortical areas is a subtype that has been consistently reported in many studies [6,7,13,21,22,33]. Interestingly, our current study disentangled the observed hippocampal sparing pattern in two different clusters with atrophy in the precuneus and the inferior parietal lobe. A unique characteristic of the most atrophied group of the two is the early onset as well as the high number of years of education, which is a proxy of cognitive reserve. This group seems to decline in cognition more rapidly than any other AD group, in agreement with the cognitive reserve hypothesis of faster disease progression in subjects with high reserve once a specific threshold has been reached [34], In contrast, the less atrophied hippocampal sparing group has a late onset in the AD symptoms, which might be the reason of the less aggressive phenotype [35].

### 3.3 Comparison between longitudinal and cross-sectional AD atrophy clusters

The subjects of this study in their majority are grouped in longitudinal clusters similar to our previously published study [13]. However, subjects from the Diffuse 1 subtype of the cross-sectional study are now distributed in more clusters because of two main reasons: 1) The Diffuse 1 cluster from the cross-sectional study is a cluster that gathered the most typical AD patterns. However, the separation from the other clusters was not very clear as discussed in that study. This cluster had the highest heterogeneity within itself and in the multidimensional scaling plot it was located between the other clusters of atrophy with more distinct patterns. 2) Importantly, the longitudinal trajectories, with help of both intercepts and slopes have disentangled the courses of the disease for the subjects that before were clustered based only in one observation in that cluster (Diffuse 1 of the cross-sectional study). In the cross-sectional study [13] we observed 4 patterns of atrophy and found 5 clusters while in the current longitudinal study we identified 3 main patterns of atrophy in 6 clusters. The existence of two different patterns of atrophy within the hippocampal sparing subtype (with differences in the AD onset) remains to be validated in larger datasets, whilst shows the potential in this method to identify them. Altogether, these findings highlight the importance of longitudinal clustering methods to advance our current ability to unravel disease heterogeneity. Our current findings show that a certain proportion of the heterogeneity may be missed by cross-sectional clustering.

There are also other aspects that differ between cross-sectional and longitudinal clustering. The statistical approach of the longitudinal clustering is based on distributional assumptions (each cluster has multivariate normal distribution), while the cross-sectional clustering was distance-based (random forest). Therefore, the longitudinal model could accommodate fixed effects (variables that we want to account for), while the cross-sectional model could not (we de-trended these effects in advance). Another important methodological difference between the two approaches is the visualization of the clusters. The crosssectional design included one more step after the clustering to compare AD groups with the sample of CU subjects in terms of ROI volumes (p-value maps). This is indeed the standard approach. Instead, the longitudinal model has an internal measure of similarity between AD groups and the CU sample, namely the fitted value maps where p-values are not calculated. We achieved a comparison between healthy aging and AD clusters without overloading our dataset with statistical comparisons. More importantly, the level of difference in actual cortical thickness or volume between two clusters of subjects (fitted value) is easier to interpret biologically and clinically than the statistical differences between clusters of subjects (p-values).

### 3.4 Limitations and strengths of the study

Our study has some limitations. The sample size is limited due to two main reasons. First, we wanted to use the results of our previous study as a ground truth for the clustering. Additionally, the exclusion criteria for CU subjects and AD patients were very strict (See material and methods), to ensure that the former group resembles a true sample of the healthy population over time, while the latter group had no missing information that can bias the interpretation of the results. This was intended to be a methodological study, although some biological interpretations are done. Hence, for the methodological part we believe our current sample size is sufficient. Yet, it is our plan to replicate our current findings in a larger sample in the future so as to investigate the generalisability of the model in the AD population. Furthermore, the variable used as time component in this study was the time from the first MRI acquisition, which helped the interpretation of the results in relation to the previous cross-sectional study, but it might limit the ability to assess if a cluster of AD subjects reflects a distinct pattern of atrophy or a stage of the disease [22]. Our study has some strengths as well. We demonstrated that incorporating longitudinal information in the clustering of imaging data is possible. The analytical framework has successfully demonstrated its ability to identify outliers with dissimilar baseline and/or atrophy progression and set them aside from the more prevalent (typical and atypical) AD patterns of atrophy. We can now apply it to different imaging modalities in order to label longitudinal data and to better understand the mechanisms underlying the aging process. The estimated model makes it possible to do two more things that were not available before: 1) to estimate future levels of atrophy for any individual subject that belongs to the clusters (prognostic value) and 2) to estimate cluster assignment of new subjects that are not included in the model training (diagnostic value).

## 4. Conclusion

In conclusion, a framework for longitudinal assessment of variability in cohorts with several neuroimaging measures was successfully tested and the results show that it can be used to understand complex processes in ageing and neurodegenerative disorders.

## 5. Material and Methods

### 5.1 Participants

We used data obtained from the Alzheimer’s disease neuroimaging initiative (ADNI), a large project launched in October 2004 in North America from Michael W. Weiner, MD. The initial goal of the ADNI 1 cohort that was used for the analysis, was to gather neuroimaging data that would help to better detect and track AD in its early stages. More specifically, positron emission tomography, MRI and other data from individuals diagnosed with AD, mild cognitive impairment (MCI) and elderly CU were collected between 2004 and 2010 from different sites of USA and Canada. The inclusion criteria for AD patients were the following: 1) to fulfil the NINCDS/ADRDA probable AD criteria, 2) a Clinical dementia rating scale (CDR) global score between 0.5 and 1, and 3) an MMSE total score between 20 and 26. The exclusion criteria for AD included: the use of psychotropic medication that could affect memory, history of significant head trauma, evidence of significant focal lesions at the screening MRI, and the existence of a significant neurological disease other than AD. For the healthy cognitively unimpaired (CU) subjects, inclusion criteria were an MMSE total score between 24 and 30 and a CDR global score equal to 0. Exclusion criteria for CU subjects comprised presence of depression, MCI or dementia. For more information on the ADNI study, see http://adni.loni.usc.edu/about/.

We included all subjects with longitudinal sMRI data and available CSF data (101 AD and 113 CU) from our previously published cross-sectional study on AD subtypes [13]. This was done to be able to compare the cross-sectional and the longitudinal clustering approach in a proper way in the same set of participants. In total 75 subjects were excluded due to bad longitudinal image quality and processing results (see below). At baseline, 94% of the AD subjects were Aβ_1-42_ positive, while only 31 CUs were included, since we wanted them all to be negative for Aβ_1-42_ and Ptau. The cut-offs for Aβ_1-42_ and Ptau used, are discussed by [36]. Moreover, the CU sample was further limited by additional inclusion criteria: 1) remain as CU subjects across all the available follow-ups and not only the ones that are used in this study (0-36 months of continuous follow-up for the 31 CU subjects), 2) have longitudinal MRI for all the time points of the analysis.

Altogether, 104 individuals were included in the final analysis, 72 AD patients (72 subjects had baseline and 12 months MRI scans, and 57 subjects had a 24 months MRI scan) and 31 CU (baseline, 12- and 24 months MRI scans). Prodromal AD subjects were not utilized in the study for two reasons: 1) adding subjects who may not develop dementia would introduce noise in the AD atrophy patterns and 2) by adding subjects who were not included in our cross sectional study, we would introduce external heterogeneity in our controlled sample and our results would not be methodologically comparable.

### 5.2 MRI acquisition and preprocessing

The MRI dataset consists of high-resolution sagittal 3D 1.5T T1-weighted Magnetization Prepared RApid Gradient Echo (MPRAGE) volumes (voxel size 1.1×1.1×1.2 mm^3^). Full brain and skull coverage were required and detailed quality control (QC) was applied to all the images [37].

Images underwent pre-processing with the longitudinal stream of the FreeSurfer pipeline (version 6.0), where a subject specific template is used [38]. Information about the FreeSurfer pipeline can be found in the following link (http://surfer.nmr.mgh.harvard.edu/fswiki/FreeSurferAnalysisPipelineOverview). Parcellation with the Desikan-Killiany [39] atlas was applied in order to extract regional average cortical thickness values. For this study we utilized cortical thickness values for 34 cortical regions and 7 subcortical volumes (hippocampus, amygdala, putamen, caudate, thalamus, accumbens, pallidum) from each hemisphere (Supplementary table 2). Estimated total intracranial volume (eTIV) was also extracted for the needs of the statistical modelling of the volumetric data [40]. This segmentation approach has previously been used for multivariate classification of Alzheimer’s disease and healthy controls, neuropsychological-, image analysis and biomarker discovery [32,41,42]. All data was processed through theHiveDB system [43]. The FreeSurfer output underwent manual visual QC to find errors in parcellations/segmentations to ensure optimal estimation of thickness and volumes. After QC, 28 AD and 48 CU subjects were excluded because of low output quality, image quality, or because less than two continuous time points existed per subject after the QC. Finally, one AD patient was excluded due to failed parcellation of regions that are included in the analysis.

### 5.3 Statistical analysis

#### 5.3.1 Data standardization

The cortical thickness and subcortical volume ROI data of AD patients were standardized based on the sample of cognitively unimpaired subjects, including mean centering and unit variance scaling. The two main benefits of mean centering and unit variance scaling of the patients’ data are: 1) after the transformation, each ROI value will represent how many standard deviations below the average CU an AD subject’s value is; 2) since we have volume and thickness data, after transformation, all the variables will have the same unit and fair statistical comparisons will be possible (without affecting the kurtosis or skewness of the distributions). This transformation has previously been applied for cross-sectional assessment of AD subtypes [8,19,44], In this study, we adapted this procedure to longitudinal data in order to account for the atrophy that is caused by the normal ageing process in the CU group over time. First we matched the timepoints of CU with the AD subjects so that the time between the initial visit of any AD subject and a longitudinal observation will match exactly with the time after the initial visit of the CU group and longitudinal observations. This will ensure that the aging time interval is accounted for. The z-values were calculated with the following formula 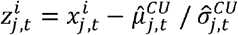, where *x* is the original measurement of subject *i*, in the time point *t* for the region *j*, while 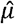 and 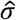 are the mean and standard deviation of the CU group at time *t* and region *j*. After this calculation, each value will resemble an atrophy level corrected for normal aging levels and also normal decline over time, which was not done previously, and is crucial for biological and clinical interpretation of brain atrophy.

#### 5.3.2 Statistical longitudinal clustering

We set out to test an analytical framework that enables us to investigate longitudinal patterns in sMRI feature analysis. For this reason, we considered a multivariate mixture model that allows us to incorporate many brain regions in the model. Moreover, in our effort to establish a general framework that will be able to facilitate both continuous and discrete data trajectories in the clustering, while accounting for the longitudinal design of the study, we decided to choose a generalized linear instead of general linear (mixed effects) approach. In addition, such an approach allows us to incorporate fixed and random effects that can serve in different ways in sMRI and other modalities. The algorithm clusters the random intercepts and slopes of each individual’s outcomes of interest (ROI measures in this study) with repeated measurements instead of repeated measurements data of each individual subject. A pair of subjects with similar estimated trajectories of atrophy (similar starting value/intercept and slope over time) will be grouped together, while subjects with different trajectories will be assigned to different groups. More specifically, as previously discussed in [45], if we know the number of clusters K, we can formulate the unobservable cluster allocation of subjects *i* as *P*(*U^i^* = *k; w*) = *w_k_, k*=1,..,K and i=1,..N. Here *w* is the vector of unknown cluster proportions that are positive and sum to 1. The meaning of *U_i_* is that *U_i_ = k* when an observation ***Y**^i^* is produced by the model density *f_i,k_*(*y_i_,p_k_,p*) where *p_k_* are cluster specific parameters and *p* are population parameters. The marginal density of *Y_i_* is 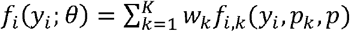, where *θ* = {*w^T^,p_k_,p*} is the vector of unknown model parameters. Finally, clustering is based on the estimated values 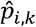 that resemble the chances of an individual *i* to belong in cluster *k* [45].

From the linear mixed model literature we know that the conditional mean response for each region *j* in a regression can be expressed as

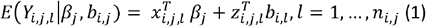

Here, *x_i,j,l_* are vectors of known covariates (fixed effects), *z_i,j,l_* is a vector that includes time values that the observations were taken, *β_r_* is the vector of unknown fixed effect for region *j, b_j_* are i.i.d. random variables that express the *j_th_* response of subject *i*. A Gaussian distribution is used to model the ROI data (general linear model), but data of ordinal or nominal nature can be analysed by changing the link function on the left part of the equation (1). For each individual i, the conditional distribution of the joint random effects vector 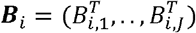 over *j* regions, given that the *i_th_* subjects belongs to the *k_th_* cluster (that is L7¿ = *k*, for a K cluster solution) follows multivariate normal distribution, is

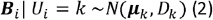

In (2), ***μ**_k_* is the unknown mean vector over *j* regions and *D_k_* is the cluster-specific positive definite covariance matrix. For each individual *i, Y_i,j,l_* (*j* = 1,…,*j* and *l* = 1,…, *n_i,j_*) are conditionally independent given the random effects ***B**_i_*. The random vectors *Y*_1_,…, *Y_N_*, as well as the random effect vectors ***B***_1_,…,***B**_N_* that were mentioned above, are independent. In summary, the *μ_k_ and D_k_*, comprise the cluster-specific parameters that we will estimate given the data to explore the various gray matter atrophy patters. The dependence among the *Y_i,j_s* (different markers *j*) is included in the non-diagonal components of the matrix *D_k_*. As mentioned in the introduction, accounting for external effects that might drive the resulting clusters within the model is convenient in this kind of analysis. Therefore, fixed effects *β_j_* in (1), common to all clusters (population level effects) are estimated for each of the external variables and brain region that we want to assess during the clustering analysis. Adding the regression dispersion parameters *ϕ_j_* to the *β_j_* vector, we summarise the model parameters common to all clusters. Those parameters are also assigned distributions. For more information specific to the distributions of those effects and their hyperparameters, please see the supplementary material (model specification).

Such an approach allows fitting the resulting cluster profiles (atrophy maps) for different combinations of fixed effects to investigate their regional contribution. Finally, since the longitudinal data from almost all cohorts with MR acquisitions typically have different numbers of visits per subject (irregularly sampled), we chose a model that can utilize all available measurements of each individual subject to calculate regression intercepts and slopes (vectors *b_i,l_*). In the model, hierarchically centered generalized LMM are assumed with a non-zero unknown mean, *b_l_* for *l* = 1,…*L* regions (see formula 1). The model that combines all the aforementioned features (Multivariate Mixture of Generalized Mixed effect Models (MMGLMM), Arnošt Komárek and Komárková 2013), is applied to longitudinal trajectories of atrophy to study whether they vary within the AD dementia spectrum (see packages “mixAK” and “coda”, R version 3.0.0 or higher). We chose a Bayesian approach for the clustering optimization because it includes prior distributions in the parameter estimates. This enables the algorithm to investigate the parameter space even in cases of small samples where likelihood information is limited.

The clustering algorithm estimates different outcomes. One outcome is the different cluster components. Each estimated multivariate Gaussian component resembles a pattern of atrophy that is observed in the dataset. Each individual subject is assigned a probability to belong to any of the components (soft clustering) rather than being assigned to a single component. The assignment of subjects into clusters is based on the maximum posterior probability rule (an individual is assigned to the component with the highest individual component probability). This is a much more realistic approach in comparison to hard clustering approaches used in most previous data-driven studies [7,8,10,17,33,46], since heterogeneity in AD is modelled here as a continuum and allows for mixed patterns instead of single patterns. Hence, the data-driven algorithm provides explicit information on whether a subject has a distinct atrophy pattern or a mixture of patterns through the estimation of subject component probabilities. The proposed framework clusters subjects of a cohort into groups (provides probability of subjects to belong in any of the clusters) and not patterns of atrophy into groups for a cohort (clusters of regions/vertices) as in the study of Marinescu and colleagues [14],

A schematic representation of the proposed analytical framework is portrayed in Figure 1. The time from the first visit (baseline) was defined as a random effect for the sake of comparability with the previous literature on AD subtypes where only one observation for each subject is included [13,17]. Therefore, the intercept of the model will correspond to the atrophy levels on the first visit and the slope will show how these atrophy levels change over the months after the first visit. The fixed effects of the model are age, sex, education, years from the onset of dementia, total intracranial volume, baseline CSF Aβ_1-42_ and pTau_181_. The resulting clusters are visualized in terms of their fitted values on the median intercept (i.e. baseline), 12 months and 24 months after the baseline observation for a specific set of fixed effects and only fitted values below 2 standard deviations of the CU mean are presented (only when values are below 95% of the CU sample) [47], Measures of dispersion (1^st^ and 3^rd^ estimate distribution quartile) are also visualized in order to assess within-cluster variance of cortical and subcortical atrophy. With those measures we can interpret how different the subjects within each cluster can be. Since this model is proposed for neuroimaging data, we also present the cortical maps of each individual and time point that was used in the analysis in the supplementary material to show how well the estimated components represent the individuals that are assigned to it.

The statistical model that we chose to employ has all the features that were described above and its original specification can be found in the supplementary material of that study [26]. The optimization was performed using the R language, version 3.4.1 [48]. The model is fully Bayesian and thus the output of the Markov chain Monte Carlo (MCMC) simulation is exploited to make inference on the population- and cluster-specific parameters. To adequately explore the distributions of the estimated parameters and speed up convergence of the algorithm, we optimized the model from different initial values based on i) the packages’ default values (see supplement to Arnošt Komárek & Komárková, 2013), ii) previous study results [13] and iii) cross-sectional clustering on the baseline data including k-means clustering and hierarchical agglomerative clustering as well as the addition of uniform noise to increase randomness in the initialization [27,49]. To identify the optimal solution for our dataset, we initially optimized models for 2-8 clusters for all the different initializations, summing to 49 MCMC chains. Then we assessed i) the model deviances (−2*logLikehood) [26], ii) the quality of parameter convergence with respect to MCMC with high autocorrelation (visual inspection of the MCMC trace plots and auto-correlation values) [27] and iii) the quality of clustering with respect to observations with low classification certainty (See Supplementary table 1 for more information). In our hybrid model evaluation approach, all three quality criteria were considered as important in the selection process (scaled to the same interval, 0-1) [50].

## 6. Author Contributions

- The study was conceived by K.P., E.W., D.F., J.B.P.
- Data were acquired, prepared, processed, or managed by K.P.
- Statistics were conceived and data were analysed by K.P., with the help from E.W., D.F., J.B.P.
- The manuscript was written by K.P. with the help from E.W.
- Data was interpreted and the manuscript was critically revised by K.P., E.W, J.B.P, D.F, Ö.S., P.V.

## 7. Acknowledgements

The authors would like to thank the Swedish Foundation for Strategic Research (SSF), The Swedish Research Council (VR), the Strategic Research Programme in Neuroscience at Karolinska Institutet (StratNeuro), the regional agreement on medical training and clinical research (ALF) between Stockholm County Council and Karolinska Institutet, Center for Innovative Medicine (CIMED), The Swedish Brain Foundation, The Swedish Alzheimer Foundation, Olle Engkvist Byggmästare Foundation, the Åke Wiberg Foundation, and Birgitta och Sten Westerberg for additional financial support, The joint research funds of KTH Royal Institute of Technology and Stockholm County Council (HMT).

Data collection and sharing for this project was funded by the Alzheimer’s Disease Neuroimaging Initiative (ADNI) (National Institutes of Health Grant U01 AG024904) and DOD ADNI (Department of Defense award number W81XWH-12-2-0012). ADNI is funded by the National Institute on Aging, the National Institute of Biomedical Imaging and Bioengineering, and through generous contributions from the following: Alzheimer’s Association; Alzheimer’s Drug Discovery Foundation; BioClinica, Inc.; Biogen Idec Inc.; Bristol-Myers Squibb Company; Eisai Inc.; Elan Pharmaceuticals, Inc.; Eli Lilly and Company; F. Hoffmann-La Roche Ltd and its affiliated company Genentech, Inc.; GE Health-care; Innogenetics, N.V.; IXICO Ltd.; Janssen Alzheimer Immuno-therapy Research & Development, LLC.; Johnson & Johnson Pharmaceutical Research & Development LLC.; Medpace, Inc.; Merck & Co., Inc.; Meso Scale Diagnostics, LLC.; NeuroRx Research; Novartis Pharmaceuticals Corporation; Pfizer Inc.; Piramal Imaging; Servier; Synarc Inc.; and Takeda Pharmaceutical Company. The Canadian Institutes of Health Research is providing funds to support ADNI clinical sites in Canada. Private sector contributions are facilitated by the Foundation for the National Institutes of Health (www.fnih.org). The grantee organization is the Northern California Institute for Research and Education, and the study is coordinated by the Alzheimer’s disease Cooperative Study at the University of California, San Diego. ADNI data are disseminated by the Laboratory for Neuro Imaging at the University of California, Los Angeles. Data used in preparation of this article were obtained from the Alzheimer’s Disease Neuroimaging Initiative (ADNI) database (adni.Ioni.ucla.edu). As such, the investigators within the ADNI contributed to the design and implementation of ADNI and/or provided data but did not participate in the analysis or writing of this report. A complete listing of ADNI investigators can be found at: http://adni.loni.ucla.edu/wp-content/uploads/how_to_apply/ADNI_Acknowledgement_List.pdf.

## Footnotes

1 Links to Europond and TADPOLE https://tadpole.grand-challenge.org/ and http://europond.eu/software/

