## Supplemental material for "Fully Bayesian longitudinal unsupervised learning for the assessment and visualization of AD heterogeneity and progression"

**Supplementary material**

|  | Number of cluster | | | | | | | | | | | | | | | | | | | | | |
| --- | --- | --- | --- | --- | --- | --- | --- | --- | --- | --- | --- | --- | --- | --- | --- | --- | --- | --- | --- | --- | --- | --- |
|  | 2 | | | 3 | | | 4 | | | 5 | | | 6 | | | 7 | | | | 8 | | |
| Simulation initial values | Criterion 1 | Criterion 2 | Criterion 3 | Criterion 1 | Criterion 2 | Criterion 3 | Criterion 1 | Criterion 2 | Criterion 3 | Criterion 1 | Criterion 2 | Criterion 3 | Criterion 1 | Criterion 2 | Criterion 3 | Criterion 1 | Criterion 2 | Criterion 3 | Criterion 1 | | Criterion 2 | Criterion 3 |
| Previous study | 26116,0 | 1 | 0,18 | 25338,6 | 1 | 0,12 | 25630,6 | 1 | 0,12 | 41803,2 | 0 | 0,03 | 25009,6 | 2 | 0,08 | 24763,4 | 6 | 0,08 | 24639,1 | | 5 | 0,08 |
| K-means | 25632,7 | 0 | 0,17 | 25585,4 | 0 | 0,14 | 25313,2 | 3 | 0,13 | 24878,3 | 2 | 0,13 | 24189,7 | 3 | 0,11 | 24269,7 | 6 | 0,09 | 24184,6 | | 4 | 0,09 |
| Hierarchical clustering | 25710,5 | 4 | 0,20 | 25692,4 | 3 | 0,17 | 25101,9 | 5 | 0,13 | 24833,1 | 1 | 0,11 | 24758,9 | 6 | 0,10 | 24406,0 | 3 | 0,09 | - | | - | - |
| Previous study (with noise) | 25984,2 | 0 | 0,18 | 25311,2 | 1 | 0,12 | 25340,3 | 5 | 0,13 | 25790,2 | 5 | 0,09 | 24983,0 | 7 | 0,08 | 24839,6 | 8 | 0,08 | 24634,5 | | 3 | 0,06 |
| K-means (with noise) | 25592,7 | 0 | 0,19 | 25577,4 | 0 | 0,17 | 25316,9 | 0 | 0,11 | 24888,2 | 0 | 0,13 | 24255,2 | 0 | 0,11 | 24312,1 | 0 | 0,09 | 24318,7 | | 0 | 0,08 |
| Hierarchical clustering (with noise) | 26288,8 | 0 | 0,22 | 25443,0 | 0 | 0,18 | 25246,4 | 0 | 0,11 | 24571,9 | 0 | 0,10 | 24473,5 | 0 | 0,10 | 24161,7 | 0 | 0,09 | 24408,9 | | 0 | 0,07 |
| Package default values | 25605,0 | 0 | 0,19 | 26352,7 | 0 | 0,08 | 47471,9 | 0 | 0,07 | 46386,7 | 0 | 0,04 | 42471,4 | 0 | 0,03 | 46843,7 | 0 | 0,15 | 46440,2 | | 0 | 0,06 |

Supplementary table 1. Model assessment

Criterion1: Deviance, criterion 2: Observations with low certainty, criterion 3: Percentage of chains with high autocorrelation. Seven different strategies were used to initialize the model and a set of three criteria were calculated from each simulation to assess the resulting model quality: 1) Deviance (the model deviance), 2) observations with low certainty (After the calculation of the highest posterior density (HPD) intervals of the individual component probabilities of a subject to belong to any cluster, those subjects for whom the lowest HPD probability was lower than 0.5 were summed. The concept behind this quality assessment method, is that if the components (clusters) estimated from the optimization process cannot express sufficiently most of the AD subjects then they are not representative of the AD subsamples and therefore they do not represent AD pathophysiology adequately), 3) Percentage of chains with high autocorrelation (the MCMC chains that were auto-correlated were summed to provide a measure of general convergence).

Supplementary table 2. List of cortical and subcortical ROIs that were included in the analysis.

| Cortical regions (thickness) | Subcortical regions (volume) |
| --- | --- |
| Banks superior temporal sulcus | Thalamus-Proper |
| Caudal anterior-cingulate cortex | Caudate |
| Caudal middle frontal gyrus | Putamen |
| Cuneus cortex | Pallidum |
| Entorhinal cortex | Hippocampus |
| Fusiform gyrus | Amygdala |
| Inferior parietal cortex | Accumbens-area |
| Inferior temporal gyrus |  |
| Isthmus–cingulate cortex |  |
| Lateral occipital cortex |  |
| Lateral orbital frontal cortex |  |
| Lingual gyrus |  |
| Medial orbital frontal cortex |  |
| Middle temporal gyrus |  |
| Parahippocampal gyrus |  |
| Paracentral lobule |  |
| Pars opercularis |  |
| Pars orbitalis |  |
| Pars triangularis |  |
| Pericalcarine cortex |  |
| Postcentral gyrus |  |
| Posterior-cingulate cortex |  |
| Precentral gyrus |  |
| Precuneus cortex |  |
| Rostral anterior cingulate cortex |  |
| Rostral middle frontal gyrus |  |
| Superior frontal gyrus |  |
| Superior parietal cortex |  |
| Superior temporal gyrus |  |
| Supramarginal gyrus |  |
| Frontal pole |  |
| Temporal pole |  |
| Transverse temporal cortex |  |
| Insula cortex |  |

Supplementary table 3. Individual images of outlier clusters and HPD uncertain group.

| RID | Group | Sex | Age | Age at the AD onset | Years of Education | APGEN 1 | APGEN 2 | CSF Abeta 42 | CSF Ptau 181 | CSF Alpha synuclein | MMSE score | MMSE Pentagon |
| --- | --- | --- | --- | --- | --- | --- | --- | --- | --- | --- | --- | --- |
| 404 | 7 | F | 88 | 84 | 14 | 3 | 3 | 234 | 32 | 1,3 | 20 | 2 |
| 404 | 7 | F | 89 | 84 | 14 | 3 | 3 | 234 | 32 | 1,3 | 22 | 1 |
| 404 | 7 | F | 90 | 84 | 14 | 3 | 3 | 234 | 32 | 1,3 | 20 | 1 |
| 1341 | 8 | F | 72 | 68 | 12 | 3 | 4 | 136 | 37 | 1,0 | 24 | 1 |
| 1341 | 8 | F | 73 | 68 | 12 | 3 | 4 | 136 | 37 | 1,0 | 22 | 1 |
| 1341 | 8 | F | 74 | 68 | 12 | 3 | 4 | 136 | 37 | 1,0 | 23 | 1 |
| 724 | 8 | M | 79 | 77 | 20 | 3 | 3 | 143 | 45 | 1,0 | 21 | 2 |
| 724 | 8 | M | 80 | 77 | 20 | 3 | 3 | 143 | 45 | 1,0 | 19 | 2 |
| 724 | 8 | M | 81 | 77 | 20 | 3 | 3 | 143 | 45 | 1,0 | 6 | 2 |
| 1281 | HPD | F | 78 | 68 | 16 | 4 | 4 | 94 | 41 | 0,7 | 25 | 2 |
| 1281 | HPD | F | 79 | 68 | 16 | 4 | 4 | 94 | 41 | 0,7 | 24 | 1 |
| 1281 | HPD | F | 80 | 68 | 16 | 4 | 4 | 94 | 41 | 0,7 | 21 | 1 |
| 753 | HPD | M | 65 | 63 | 16 | 4 | 4 | 129 | 62 | 4,2 | 24 | 1 |
| 753 | HPD | M | 66 | 63 | 16 | 4 | 4 | 129 | 62 | 4,2 | 20 | 1 |
| 753 | HPD | M | 67 | 63 | 16 | 4 | 4 | 129 | 62 | 4,2 | 17 | 1 |
| 852 | HPD | F | 84 | 83 | 18 | 3 | 4 | 131 | 32 | NA | 24 | 1 |
| 852 | HPD | F | 85 | 83 | 18 | 3 | 4 | 131 | 32 | NA | 22 | 1 |

The variable group here has three different values: 1 = Cluster 1, 7= cluster 7, HPD = High posterior density interval uncertain classification. APGEN 1 and 2 refer to the Apoe E4 alleles (3 is an Apoe E3 carrier and 4 is and Apoe E4 carrier). CSF values are in pg/ml.

Supplementary figure 1. 1^st^ and 3^rd^ quartile fitted value cortical thickness images

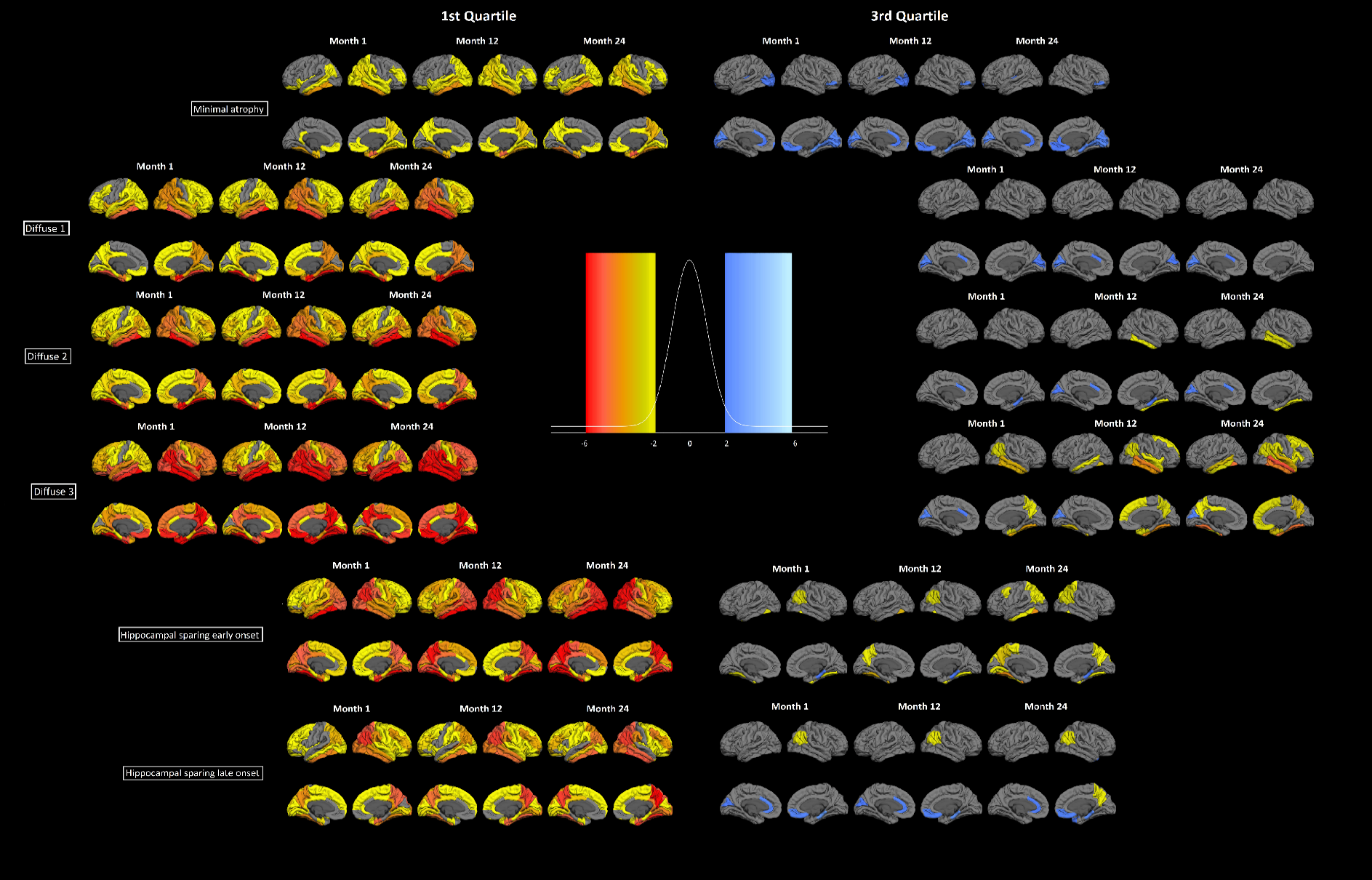

Fitted values of the 6 longitudinal atrophy clusters for the AD sample. Each row presents the quartile 1 and quartile 3 fitted value images of the cortical and subcortical atrophy of the 6 clusters for three time point (1, 12 and 24 months from the first measurement). The data are presented as controls z-scores. Fixed effects: Intracranial volume = average Intracranial volume, Sex= female, Age = 75 years, AD duration = 5 years, Education = 16 years, CSF Abeta 42 = 100 pg/ml, CSF Ptau181P = 50 pg/ml. These images help to understand and characterize each cluster separately.

Supplementary figure 2. Individual cortical atrophy pattern for each cluster’s input data.

A) Minimal atrophy

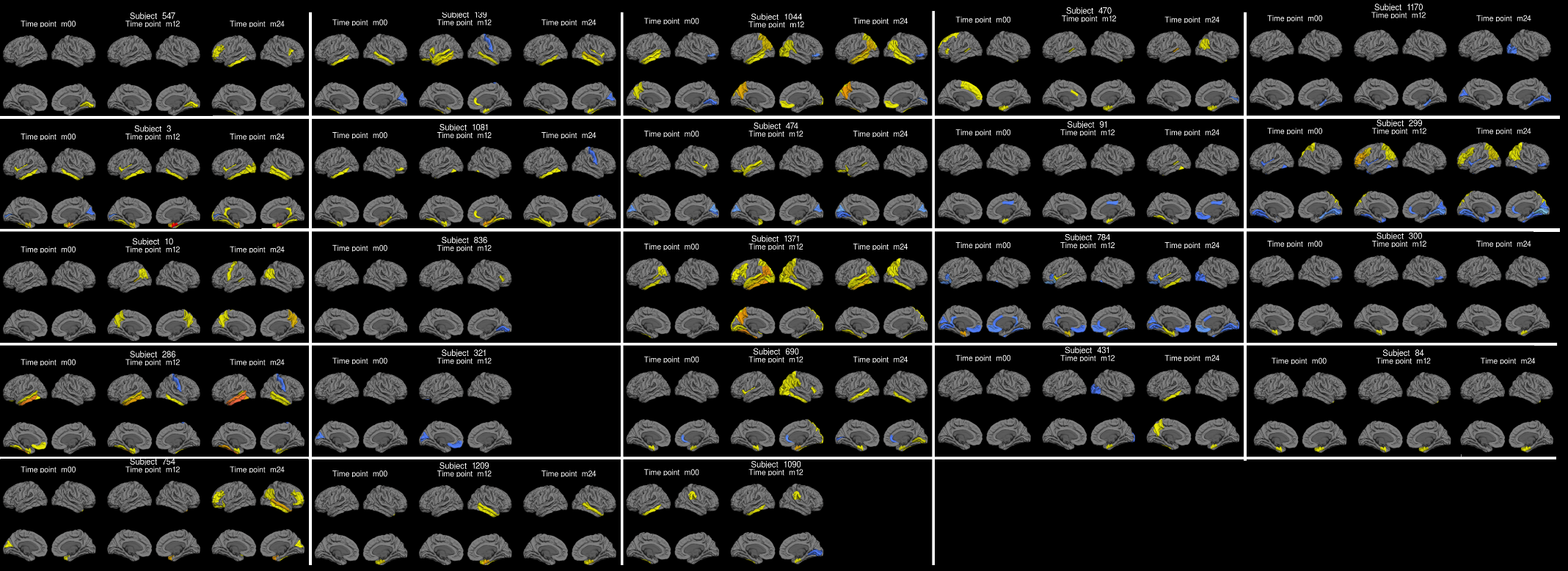

B) Diffuse 1

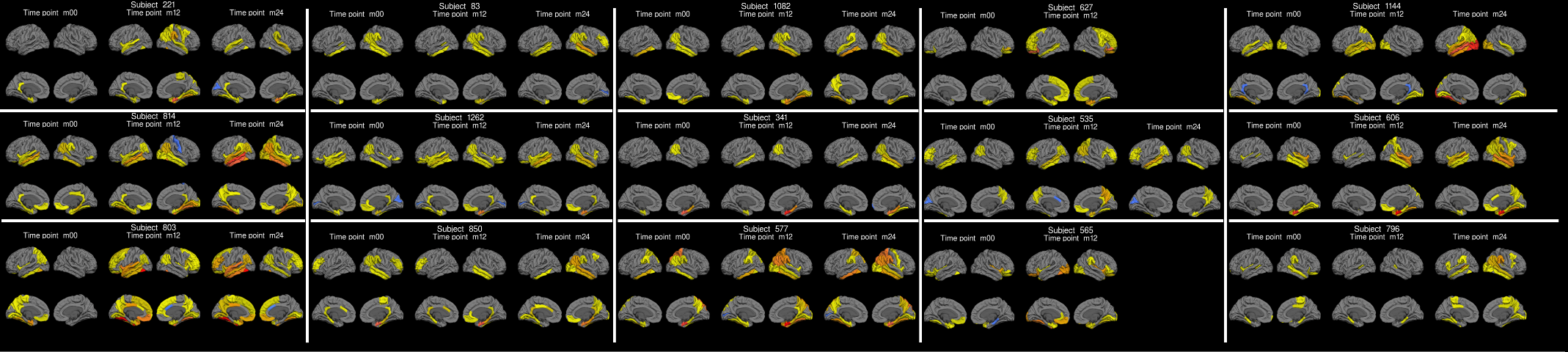

C) Diffuse 2

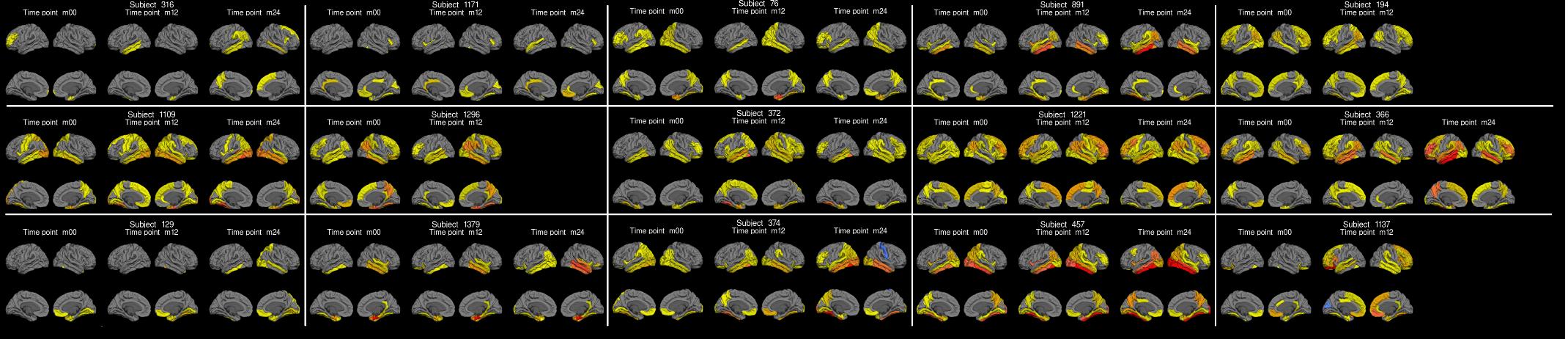

D) Diffuse 3

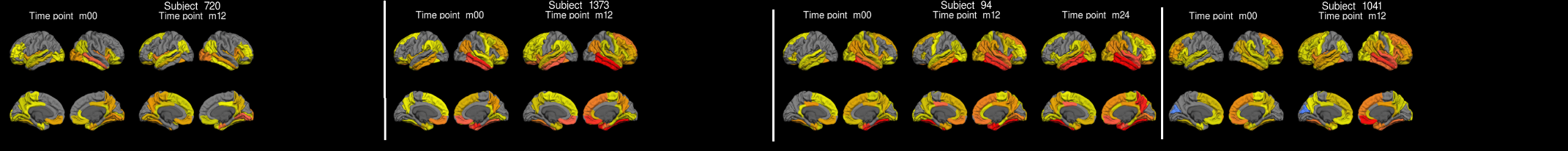

E) Hippocampal sparing early onset

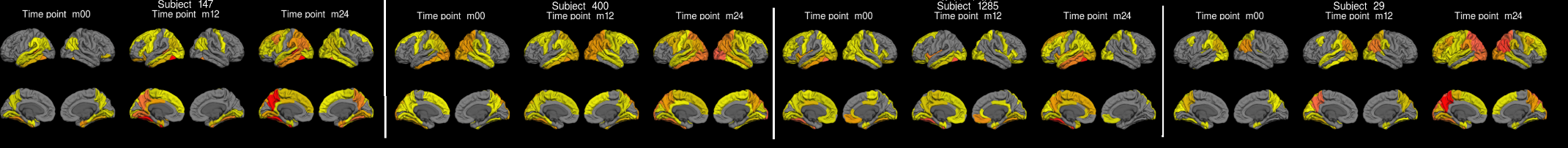

F) Hippocampal sparing late onset

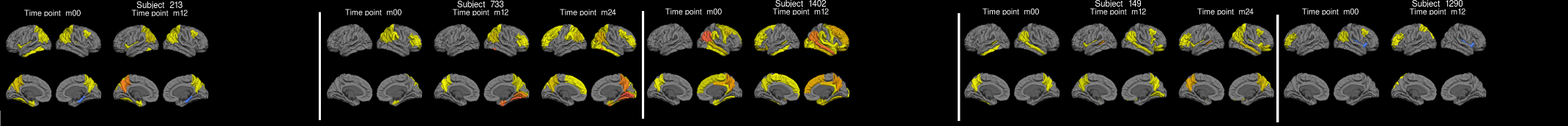

This is a visualization to increase the transparency of the clustering procedure. Each cell (top row: subject ID) comprises the cortical thickness of one subject for their different time points (second row, Time point m00: baseline observation, Time point m12: 12 months follow up, Time point 24: 24 months follow up). The subject atrophy maps are presented in terms of z-values after a linear correction for the same effect variables as the clustering algorithm fixed effects. The legend of colours is the same as in the supplementary figure 1. Only thickness that exceeds 2 standard deviations from the control group regional distributions are presented. A) Minimal atrophy B) Diffuse 1, C) Diffuse 2, D) Diffuse 3, E) Hippocampal sparing early onset, F) Hippocampal sparing late onset.

This visualization helps to understand what the data that the algorithm has as input look like. This way it is easier to understand why the resulting clusters have the component patterns that are observed in figure 3 of the manuscript.

Supplementary figure 3. Demographics of outlier clusters and HPD uncertain group.

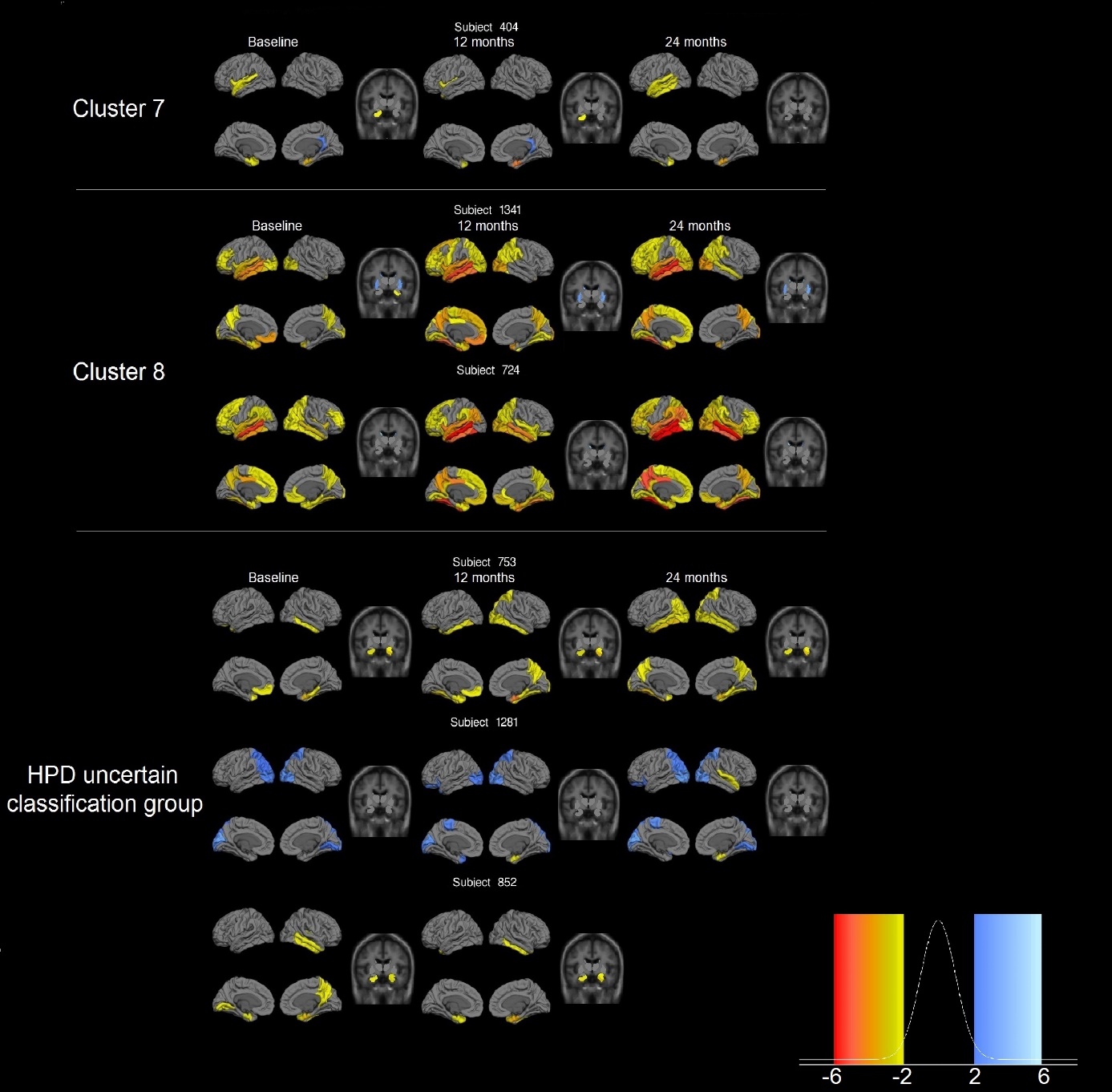

The individual subject images have the same colour legend as in supplementary figure 1 and therefore are presented in terms of standard deviations above or below the average of the control group. The outlier cluster 7 consists of one subject, the cluster 8 of 2 subjects and the HPD uncertain classification group of 3 subjects. The cluster 1 and cluster 7 are outlier clusters, while the HPD group subjects are subjects that could not be classified with high certainty in any of the 6 main clusters of the analysis result.

Supplementary figure 4. Trajectories of MMSE scores over time.

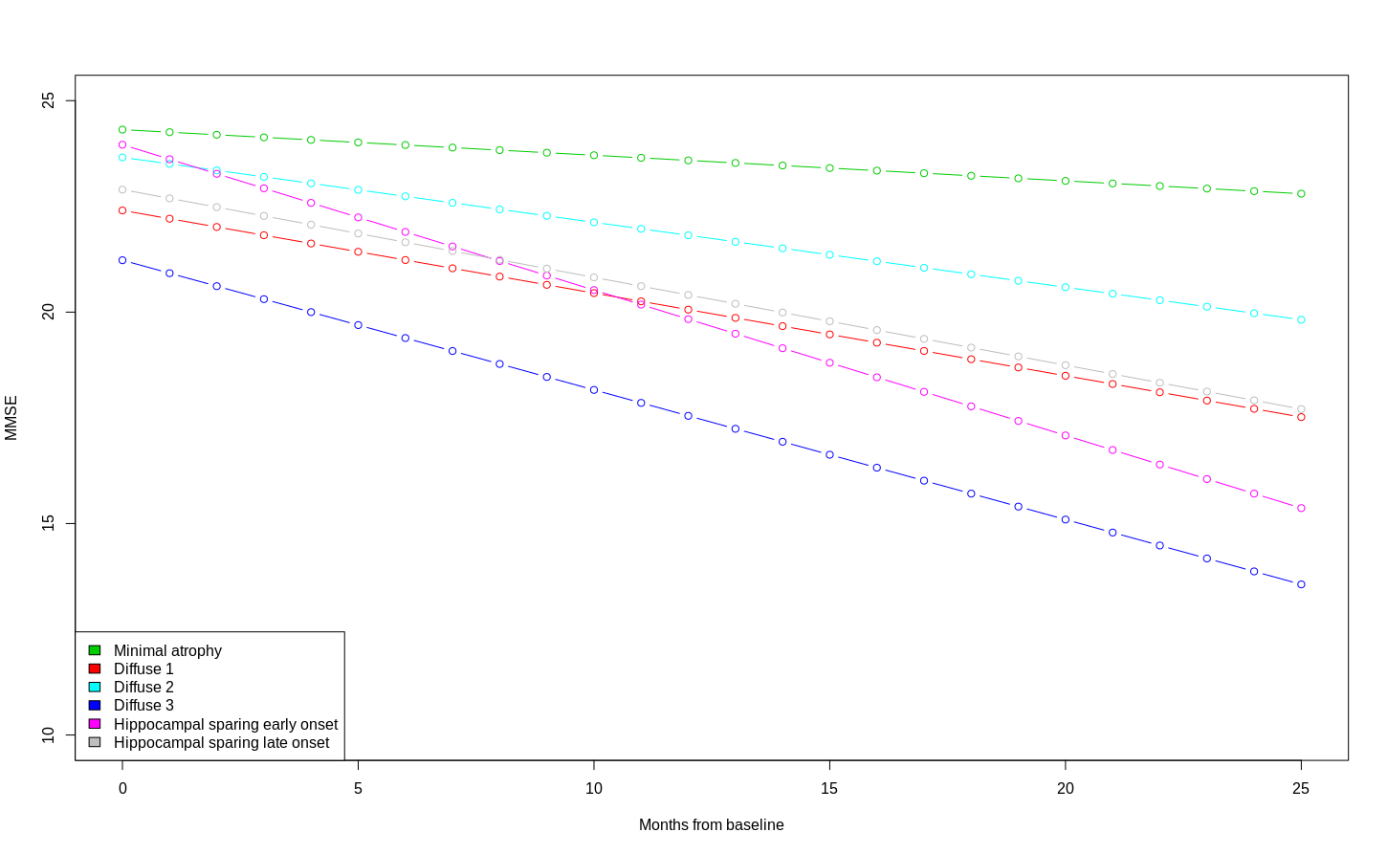

The trajectories of decline in total MMSE score were calculated for the six clusters. Mixed effect linear regression was employed, and the model followed all the assumptions for linear models. The fixed effects were used to assess the differences at baseline and in decline between the 6 groups presented at the main part of the manuscript. The random effects included the subject ids to account for dependencies in the repeated measurements. Significance of fixed effects was not assessed, and the interpretation was based in the coefficient standard errors. The minimal atrophy cluster had the highest baseline MMSE and maintained it during the course of the available assessments. The diffuse 3 group had the lowest baseline scores and a steep decline in MMSE. However, the steepest decline in MMSE was observed in the hippocampal sparing early onset cluster that started at levels comparable to the minimal atrophy cluster and reached the levels of diffuse 3 cluster after 25 months.

Supplementary material: model specification

- Cluster-specific effect

The normally distributed random effects $\beta_{i,l}$have parameters$\mu_{k}$ and$D_{k}$. The random parameter $\mu_{k}$ follows normal distribution with mean 0 and positive definite diagonal covariance matrix with 6 in the diagonal and 0 in the non-diagonal elements imposing independent apriori means. The inverted mixture covariance matrix $D_{k}^{-1}$ follows independent Wishart distributions with 165 degrees of freedom and the diagonal covariance matrix with elements random parameters $\gamma$ that follow gamma distribution with hyperparameters for shape = 0.2 and scale = 0.3. The a priori distribution for the proportions $w_{k}$ is Dirichlet with parameters equal to 1.

- Population-specific effects
  - The population parameters $\beta_{j}$follow normal distribution with hyperparameter mean 0 and standard deviation 100. This is a relatively uninformative prior and we checked that the posterior standard deviations were much lower after being estimated with our data.
  - The regression dispersion parameters followed a gamma distribution (dispersion parameters often have either gamma or Wishart distribution since those distributions take only positive numbers, that is support over 0) with parameters for shape equal to 1 and the scale was a random hyperprior. The inverted hyperprior follows a gamma distribution too, with shape equal to 0.2 and scale selected by the routines of the package mixAK in R given our data.
